# Challenges and perspectives of quantitative functional sodium imaging (fNaI)

**DOI:** 10.1101/375055

**Authors:** Claudia AM Gandini Wheeler-Kingshott, Frank Reimer, Fulvia Palesi, Antonio Ricciardi, Gloria Castellazzi, Xavier Golay, Ferran Prados, Bhavana Solanky, Egidio U D’Angelo

## Abstract

Brain function has been investigated via the blood oxygenation level dependent (BOLD) effect using magnetic resonance imaging (MRI) for the past decades. Advances in sodium imaging offer the unique chance to access signal changes directly linked to sodium ions (23Na) flux across the cell membrane, which generates action potentials, hence signal transmission in the brain. During this process 23Na transiently accumulates in the intracellular space. Here we show that quantitative functional sodium imaging (fNaI) at 3T is potentially sensitive to 23Na concentration changes during finger tapping, which can be quantified in grey and white matter regions key to motor function. For the first time, we measured a 23Na concentration change of 0.54 mmol/l in the ipsilateral cerebellum, 0.46 mmol/l in the contralateral primary motor cortex, 0.27 mmol/l in the corpus callosum and −11 mmol/l in the ipsilateral primary motor cortex, suggesting that fNaI is sensitive to distributed functional alterations. Open issues persist on the role of the glymphatic system in maintaining 23Na homeostasis, the role of excitation and inhibition as well as volume distributions during neuronal activity. Hemodynamic and physiological signal recordings coupled to realistic models of tissue function will be critical to understand the mechanisms of such changes and contribute to meeting the overarching challenge of measuring neuronal activity *in vivo*.

## Introduction

Ever since the blood oxygenation level dependent (BOLD) effect was described, functional magnetic resonance imaging (fMRI) has dominated neuroscience as a mean to evaluate brain activity (Ogawa et al., 1990b, 1990a, 1992). It allows mapping of signal changes generated by the mismatch between oxygen delivery and consumption upon neuronal activation. While providing significant insight into brain function, a major limitation of BOLD is that it is an indirect measure of function and is affected by subject-specific haemodynamic factors. Thus, an approach that could directly measure neuronal activity in humans *in vivo*, non-invasively, would have a major advantage over BOLD-fMRI.

In this paper, we propose that, thanks to sodium MRI technology, we are within reach of measuring directly a local transient change of sodium ions (23Na) concentration in the intracellular space. During activity, neuronal action potentials cause a transient 23Na flux from the extracellular to intracellular space over a temporal scale of several milliseconds. If sodium imaging, dynamically repeated, was successful in detecting 23Na concentration changes evoked by specific tasks, it would open up a new way of investigating human brain function, complementing BOLD-fMRI.

Imaging aspects of the brain electrical activity, other than BOLD, related to transmembrane sodium-potassium ion exchange during depolarization could provide a direct access to primary brain function everywhere. Although sodium channels are predominantly located in the axonal initial segment (Chadderton et al., 2004; Dover et al., 2016; Häusser and Clark, 1997; Masoli et al., 2015; Powell et al., 2015; Rancz et al., 2007), they are also expressed in Ranvier nodes along the white matter (WM) axons. While BOLD signals capture mainly the large energy demand supporting brain function related to the sodium-potassium pump to re-establish ionic gradients after action potentials in gray matter (GM) (Brockhaus et al., 1993; Koch and Barish, 1994), 23Na concentration changes could be sensitive to activity also in WM and therefore contribute to our understanding of brain circuits involved in specific tasks.

Despite its limitations, BOLD-fMRI has been very successful in neurological research applications to study mechanisms of disease. Pathologies where blood perfusion is impaired, such as multiple sclerosis (Paling et al., 2013) and stroke (Sakatani et al., 2007), reveal alterations during task BOLD fMRI. However, these may be mediated by a dysfunction in evoked blood oxygenation or by neuronal damage itself. Experimental neurophysiology also indicates that psychiatric conditions, such as autism, are characterised by altered patterns of neuronal firing, which are difficult to capture *in vivo* using BOLD-fMRI (Giza et al., 2010; Leblond et al., 2014). Considering its substantial research output, BOLD-fMRI is rarely used clinically, though, besides pre-surgical planning. Yet, while growing evidence supports that minimising residual tumour mass improves survival, false functional localisation may render it less effective (Morrison et al., 2016; Suchorska et al., 2016).

This means that there is a pressing need for tools able to directly map brain function, rather than through hemodynamic effects, and with greater reliability. Again, measuring 23Na concentration changes could meet this need, complementing BOLD-fMRI, with the potential of impacting clinical practice.

From a physiological point of view, neuronal cells’ function has recently been mapped *in vitro* with high specificity, describing distribution and functionality of 23Na channels with incredible details (Dover et al., 2016; Masoli et al., 2015). In parallel, physiological studies have also led to a better understanding of the neurovascular coupling (Howarth et al., 2009; Lippert et al., 2010; Mapelli et al., 2016) at the origin of the BOLD-fMRI signal (Ogawa et al., 1990b, 1990a, 1992). These data are the bases for constructing emerging realistic models of neuronal activity, built on ever accurate physiological recordings of cellular function, and could provide an invaluable tool to interpret large scale measures of brain function from MRI (D’Angelo and Wheeler-Kingshott, 2017; Friston et al., 2017; Blanchard et al., 2016).

From a technological point of view, it is now feasible to measure 23Na concentrations *in vivo*, which are key in retaining physiologically balanced tissues. Indeed, it is now possible to non-invasively measure quantitatively total (i.e. intra + extra cellular) 23Na concentrations (TSC) of the human brain tissue *in vivo* using high field MRI scanners (Thulborn, 2018). Furthermore, 23Na in the intra and extracellular spaces have different MR properties due to their cellular environment, hence any alteration in volume fractions or in intra or extracellular 23Na concentrations could affect the measured TSC. Arguably one could say that TSC is sensitive to changes due to tissue composition as well as to pathological changes of cellularity, albeit with a lower sensitivity than proton (1H), or indeed due to transient TSC changes (ΔTSC) during functional activity.

With preliminary data and biophysical hypothesis of ΔTSC changes in tissue *in vivo*, we intend to establish a framework for developing functional sodium imaging (fNaI), demonstrating an exciting opportunity for measuring brain function and potentially neuronal activity, addressing a pressing need for a multi-disciplinary integration.

## Methods

### Subjects

8 right-handed healthy volunteers (mean age 33yrs, range 27-45, 5 males) gave written consent to this study approved by the NRES Committee London – Harrow, in accordance with the Declaration of Helsinki.

### fNaI acquisition protocol

Data was acquired on a 3T Philips Achieva system (Philips, Netherlands) with a single-tuned volume head-coil (Rapid, Germany) using a 4-times undersampled 3D-Cones ultra-short echo time sequence (Gurney et al., 2006; Riemer et al., 2014), 4mm isotropic resolution, 240mm field-of-view, 90° flip-angle, TR=50ms, TE=0.22ms, 6 NEX, total scan time per volume=60s. TE was defined as from the end of the pulse to the start of readout (0.22 ms). The RF pulse was 320 μs, so the time from the centre of the RF pulse to the start of readout is 0.38ms. The length of the readout was 30ms.

### fNaI paradigm design

fNaI was performed back-to-back 6-times (3-rest conditions interleaved with 3-tasks). Subjects were asked (verbally) to perform a right-hand finger-tapping task (self-pacing at a frequency of 1Hz), opposing the thumb to each one of the fingers, repeatedly from the index to the little finger and back, with ample extension of the movements.

### fNaI data analysis

Images were reconstructed to 2mm isotropic resolution using SNR-enhancing sub-Nyquist k-space sample weighting (Pipe, 2000). All analyses were performed with SPM8. Images were rigidly registered, smoothed with a 8×8×8mm^3^ Gaussian-kernel and normalised to the proton density (PD) MNI152 template. Statistical analysis was performed using the SPM8-PET group analysis toolbox. Statistical maps were calculated with p=0.001, cluster extent of k = 20 voxels and family-wise error (FWE) correction.

### fNaI clusters identification

Maps of t-statistics were saved from the fNaI data analysis and imported in the xjview toolbox (http://www.alivelearn.net/xjview) of SPM for a detailed cluster report in terms of peak, number of voxels, location and anatomical areas involved in Tailarach atlas space (Yoon et al., 2012).

### Sodium ions (23Na) flux and ΔTSC

Voxel-wise TSC was calculated according to (Christensen et al., 1996), using two reference phantoms (33 and 66 mmol/l sodium agar) placed either side of the brain for 4 out of 8 volunteers. ΔTSC was calculated from TSC on/off maps for cerebellar, ipsi and contralateral primary motor cortex (M1) and corpus callosum (CC) clusters and reported as (mean ∆TSC ± standard deviation) across the 4 subjects.

## Results

fNaI was successfully performed in 8 subjects during a right-hand finger-tapping task at 3T. Figure 1a) shows transverse slices from one fNaI volume of a randomly chosen subject, while Figure 1b) shows fNaI statistical activation maps from the group analysis. A total of 16 main clusters were identified and are reported in Table 1. These include the contralateral M1 (Precentral), somatosensory (Postcentral) and supplementary motor (Superior Frontal) gyri, and the ipsilateral anterior cerebellum (lobule I-IV) as well as lobule VI, Crus I-II and the dentate nucleus (Figure 1c)). Noticeable are the large number of ipsilateral areas that were also activated in the ipsilateral cerebrum (e.g., frontal and temporal lobes, postcentral gyrus, deep grey matter including the right thalamus, the insular cortex, the limbic lobe and parahippocampal gyrus. The lingual gyrus, Brodmann areas (BA) 2, 19 and 37 were also activated ipsilaterally to the movement). Interestingly, WM areas were identified in the CC, contralateral paracentral lobule and medial frontal gyrus, corticospinal tract (CST), posterior cingulum and ipsilateral supramarginal gyrus.

**Figure 1.**
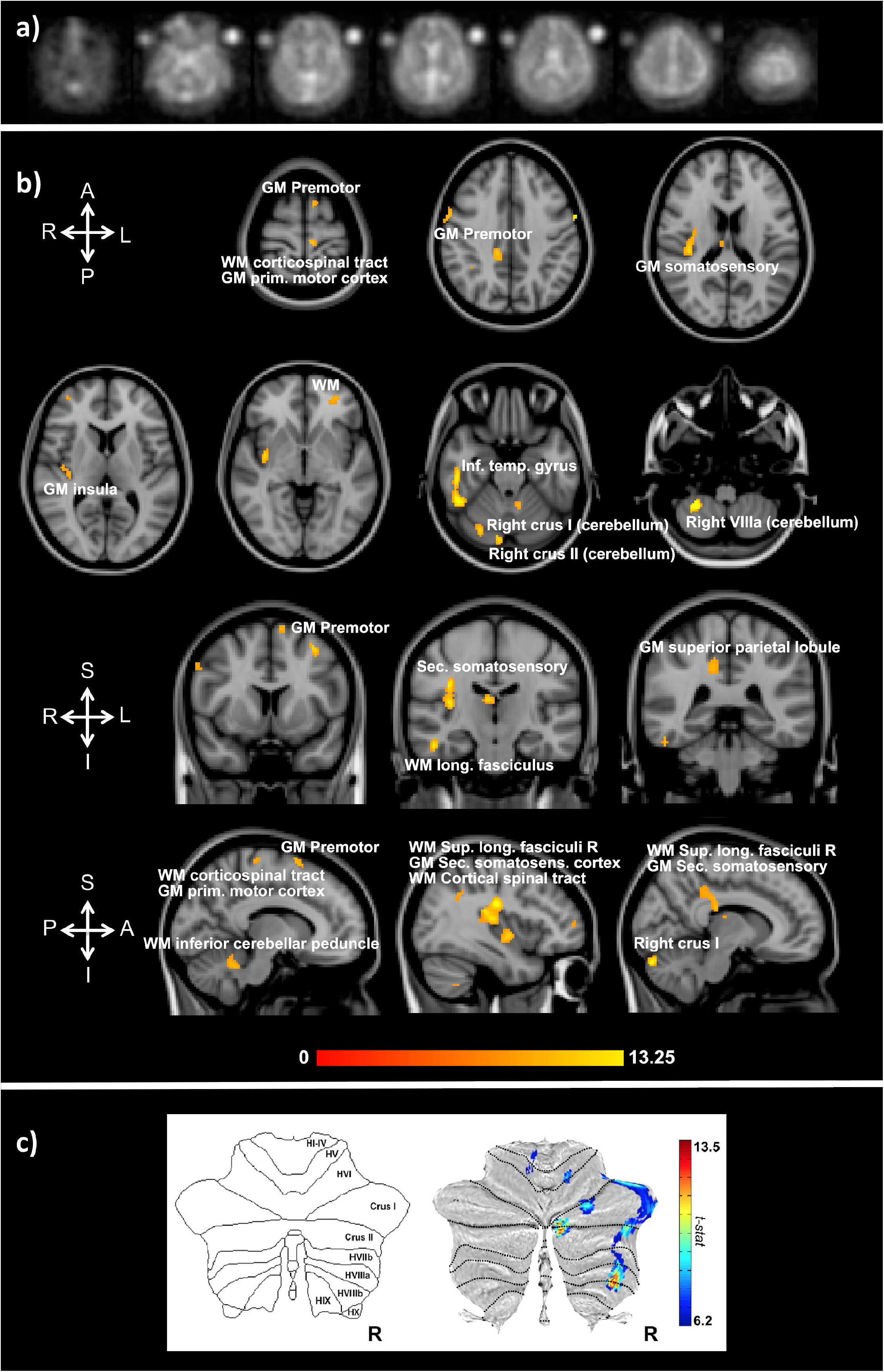
a) Example of transverse slices from a single functional sodium imaging (fNaI) volume after smoothing. The circles either side of the brain are known concentration phantoms. Signal to noise ratio in WM was measured as (17.5 ± 1.4) a.u. in all 8 subjects. b) Activation clusters from proof of concept fNaI experiment where the subject performed a 1Hz finger-tapping paradigm. Results are from group analysis of 8 volunteers (FWE corrected, p<0.001, 20 voxels), overlaid on 3D T_1_-weighted structural images with anatomical annotations. Maps were poorly localized at p<0.05, hence the higher than usual threshold. Worth noticing that signal changes for fNaI were of the order of 10%, which is twice what is normally detected using BOLD-fMRI. Activations are seen in motor-function related areas. c) The cerebellum shows enhanced activations in Crus I/II and lobule VI related to finger tapping and motor planning. GM: gray matter and WM: white matter.

**Table 1:** Clusters of activations and identification of areas involved according to the Tailarach atlas in XJVIEW. In red are right (ipsilateral) clusters and in blue are left (contralateral) clusters with respect to the hand used for the task. Total sodium concentration (TSC) values and their standard errors (SE) for each cluster across 4 of the 8 volunteers are also reported. BA = Brodmann Area

| Cluster | N voxels | Peak MNI coordinates |  |  | Peak Description | Sub clusters | Mean TSC mmol/l (SE mmol/l) |
| --- | --- | --- | --- | --- | --- | --- | --- |
|  |  | X | Y | Z |  |  |  |
| 1 | 612 | 28 | -50 | -52 | Right Cerebellum // Cerebellum Posterior Lobe // Cerebellar Tonsil // Cerebellum_8_R (aal) | Cerebellar Tonsil<br>Cerebellum 8 R (aal)<br>Inferior Temporal Gyrus BA 37<br>Fusiform R (aal)<br>Cerebellum Crus1 R (aal)<br>Middle Temporal Gyrus | 39 (4) |
| 2 | 55 | 12 | -86 | -26 | Right Cerebellum // Cerebellum Posterior Lobe // Declive // Cerebellum_Crus1_R (aal) | Declive<br>Cerebellum Crus1 R (aal)<br>Cerebellum Crus2 R (aal) | 42 (5) |
| 4 | 45 | 28 | -74 | -22 | Right Cerebrum // Cerebellum Posterior Lobe // Declive | Declive<br>Cerebellum 6 R (aal) | 36 (4) |
| 3 | 101 | -8 | -50 | -28 | Left Cerebellum // Cerebellum Anterior Lobe // Fastigium | Culmen<br>Cerebellum 4-5 L (aal)<br>Fastigium<br>Dentate | 35 (3) |
| 5 | 55 | 18 | -60 | -10 | Right Cerebrum // Occipital Lobe // Lingual Gyrus // White Matter // Lingual_R (aal) | Lingual R (aal)<br>BA 19<br>White Matter<br>Limbic Lobe<br>Parahippocampal Gyrus<br>Culmen<br>Right Cerebellum<br>Cerebellum Anterior Lobe | 42 (3) |
| 6 | 66 | -36 | 48 | -10 | Left Cerebrum // Frontal Lobe // Middle Frontal Gyrus // White Matter // Frontal_Mid_Orb_L (aal) | Frontal Mid Orb L (aal)<br>Gray Matter<br>BA 11<br>Frontal Sup Orb L (aal) | 39 (5) |
| 7 | 432 | 38 | -18 | 26 | Right Cerebrum // Sub-lobar // Extra-Nuclear // White Matter | White Matter<br>Insula R (aal)<br>BA 13<br>Extra-Nuclear<br>Rolandic Oper R (aal)<br>Heschl R (aal)<br>Frontal Lobe<br>Putamen R (aal) | 38 (3) |
| 8 | 20 | 44 | 52 | 4 | Right Cerebrum // Frontal Lobe // Inferior Frontal Gyrus // White Matter // Frontal_Mid_R (aal) | Frontal Mid R (aal)<br>Right cerebrum<br>White Matter | 30 (5) |
| 9 | 296 | 6 | -18 | 14 | Right Cerebrum // Sub-lobar // Thalamus // Gray Matter // Thalamus_R (aal) | White Matter<br>Limbic Lobe<br>Cingulate Gyrus<br>Cingulum_Mid_R (aal)<br>Sub-lobar<br>Gray Matter<br>Extra-Nuclear<br>Thalamus_R (aal)<br>Corpus Callosum<br>Frontal Lobe | 59 (6) |
| 10 | 22 | 36 | -48 | 36 | Right Cerebrum // | Supramarginal Gyrus | 33 (4) |
|  |  |  |  |  | Parietal Lobe // White Matter |  |  |
| 11 | 32 | -60 | 0 | 36 | Left Cerebrum // Frontal Lobe // Precentral Gyrus // BA 6 // Precentral_L (aal) | BA 6<br>Precentral_L (aal)<br>BA 4 | 31 (5) |
| 12 | 64 | 60 | 2 | 38 | Right Cerebrum // Frontal Lobe // precentral Gyrus // Grey Matter // BA 6 // precentral_R (aal) | Precentral Gyrus<br>BA 6<br>Inferior Frontal Gyrus<br>Postcentral R (aal) | 29 (6) |
| 13 | 26 | -34 | 8 | 52 | Left Cerebrum // Frontal Lobe // Middle Frontal Gyrus // Gray Matter // BA 6 // Frontal_Mid_L (aal) | Frontal Mid L (aal)<br>White Matter<br>BA 6 | 41 (4) |
| 14 | 24 | -46 | -32 | 58 | Left Cerebrum // Parietal Lobe // Postcentral Gyrus // Gray Matter // BA 2 // Postcentral_L (aal) | BA 2<br>Postcentral_L (aal)<br>BA 4 | 36 (4) |
| 15 | 28 | -8 | 8 | 68 | Left Cerebrum // Frontal Lobe // Superior Frontal Gyrus // White Matter // Supp_Motor_Area_L (aal) | Superior Frontal Gyrus<br>Supp Motor Area L (aal)<br>White Matter | 45 (4) |
| 16 | 25 | -8 | -28 | 68 | Left Cerebrum // Frontal Lobe // Medial Frontal Gyrus // White Matter // Paracentral_Lobule_L (aal) | Paracentral Lob. L (aal)<br>White Matter<br>Medial Frontal Gyrus | 45 (5) |

∆TSC in the anterior cerebellar cluster was (0.54 ± 0.17) mmol/l, while it was (0.27 ± 0.08) mmol/l in the CC, (0.46 ± 0.10) mmol/l in the contralateral M1; a negative ΔTSC change was measured in the ipsilateral M1 (−0.11 ± 0.06) mmol/l.

## Discussion

We have shown preliminary evidence that brain function can be assessed non-invasively by sodium imaging *in vivo* using a 3T MRI scanner, a major step forward in the overarching aim to directly assess neuronal activity. Indeed, fNaI successfully detected changes in activation between finger tapping and rest across the entire brain. Activations in motor and executive control areas indicate that fNaI has the potential to be an effective biomarker of functional activity.

The localisation of activated regions is conveying interesting results. Several clusters are at the border between GM and WM, where a high concentration of sodium channels in the axon initial segment could lead to a large intracellular 23Na accumulation (Dover et al., 2016). The task employed in this initial experiment was indeed demanding: one minute of self-paced finger taping requires motor planning and concentration. Interestingly, as well as M1 (BA4), the study shows activations in the primary somatosensory cortex (BA2), i.e. the main cortical area for processing the sense of touch, and in the premotor cortex (BA6), which is involved (with the cerebellum) in self-pacing finger tapping (Mak et al., 2016; Witt et al., 2008). In the cerebellum, activations occurred in posterior ipsilateral areas, namely Crus I/II, lobule VI, and lobule VIIIa, which are heavily involved in integrative aspects of motor control and cognitive processing (Mak et al., 2016; Witt et al., 2008). A further finding is the presence of patches of activity in WM, including: the CST, which is the major pathway for the motor system; the superior longitudinal fasciculus that connects parietal to prefrontal cortices with associative fibres integrating body awareness and perception; the CC, which connects both hemispheres; the cingulum, which receives afferent fibres from the thalamus, as part of the spino-thalamic tract. These tracts are myelinated and enriched in sodium channels at the nodes of Ranvier, forming the axonal pathways wiring-up the sensorimotor network. Given that ionic fluxes (including those involving astrocytes) are generally smaller in WM than GM, it will be important to verify these activations in future studies, to exclude possible partial volume effects and to assess whether neurotransmitter signaling could cause enough accumulation of 23Na at synapsis junctions to be detectable with fNaI. If these results were going to be confirmed, reconstruction of axonal circuits supporting functions would find invaluable information in WM matter fNaI results.

From the present data it is impossible to determine the mechanisms underlying fNaI changes, i.e. does ∆TSC reflect sensitivity mainly to the shift of 23Na between the intracellular and extracellular compartments (Gilles et al., 2017), or does it reflect also changes in vascular and perivascular spaces through neurovascular coupling between glucose metabolism and increase blood delivery? In other words, we cannot exclude that in the present measurements, a significant contribution comes from the blood. Moreover, given the experimental TR, T1-weighting could play a role in the contrast: any excess of blood-related signal flowing through the vessels in activated regions could lead to artefactual increases in sodium signal. Future fNaI studies should consider minimizing this possible effect.

Whilst a precise estimate would require sophisticated models and combinations of experimental measurements *in vivo* and *in vitro*, here we can probe likely/expected scenarios and propose some *ab initio* calculations. Can the molar flux of 23Na displaced during activity be sufficient to generate a meaningful fNaI signal? For example, in the cerebellum, ∆TSC was 0.54 ± 0.17 mmol/l. The measured ATP consumption during activity in the cerebellar cortex is 20.5 mmol of ATP/(g∙min) (Howarth et al., 2009, 2012; Sokoloff et al., 1977). Of this ATP, ~50% is used for computation while the other ~50% for maintenance (Brockhaus et al., 1993; Koch and Barish, 1994; Howarth et al., 2009, 2012), so about 10 mmol of ATP/(g∙min) are used for function. One ATP corresponds to shifting three 23Na (previously accumulated inside the cell) through the cell membrane to re-establish ionic balance; this means that during activity there is a shift of the order of 30 mmol/(g∙min) of 23Na, i.e. 30*10^3^mmol/(l∙min) of 23Na, which is orders of magnitude larger than our measured value. Therefore, there is sufficient 23Na displacement to possibly explain the fNaI signal. The larger 23Na displacement expected from calculations compared to ∆TSC is likely to reflect the fact that, while 23Na enters through sodium channels during the action potentials, soon thereafter it leaves the cell through sodium-potassium pumps and sodium exchangers. What is established during a given time-frame is a dynamic equilibrium between 23Na influx and efflux with a residual unbalance, that is potentially captured by the measured ∆TSC. But there are yet other sources of 23Na flux that should be considered. Other fluxes that contribute to reaching equilibrium are due to 23Na co-transport with other ions, metabolites and neurotransmitters in neurons and glial cells, in support of the energy budget during activation (Dienel et al., 2008; Hertz et al., 2015; DiNuzzo et al., 2017). How all of these contribute to ΔTSC sign and magnitude remains to be discovered. Furthermore, with the current 4×4×4mm^3^ resolution of fNaI, it is also possible that this displacement – which happens on a microstructure scale - is actually diluted or even averaged out. It is also important to assess the contribution of changes in cerebral blood volume (CBV) during brain function and changes in Virchow-Robin space volume (VRSV) and the glymphatic system. During functional activity CBV changes because of the arteriole (CBVa), where CBVa at rest is of the order of 0.8ml/100g, with a change of ∆CBVa=0.34ml/100g (Hua et al., 2011; Kim et al., 2007; Lee et al., 2001). This is transferred to the capillaries, where the venous side passively follows the arterioles dilation and resistance changes, which induce blood flow and volume changes (Buxton, 2012). If ∆CBVa is added to the extracellular space compartment, reducing the cellular volume fraction even by as little as 0.5%, this would be enough to cause ΔTSC of the order of the measured one, given the 10 times higher extracellular molar concentration. This ∆CBVa, though, would not affect the ΔTSC measured if its fractional volume was balanced by a corresponding reduction in VRSV, i.e. if the extracellular space (CBV + extracellular matrix + VRSV) and the cellular space (e.g. intracellular space + myelin) proportions remained constant. In this hypothesis, the perivascular space and therefore the glymphatic system, would work as a compensatory chamber, with “rigid” neuronal structures within the extracellular matrix (Thulborn, 2018). The cerebrospinal fluid role in buffering the changes in CBV during functional activity has been investigated and reported to be reduced during activation, using relaxometry in the visual cortex, which would support this hypothesis (Piechnik et al., 2009).

On the other hand, we must not forget that the ΔTSC measured experimentally in this study *in vivo* is an “apparent” TSC change, as the current acquisition protocol cannot distinguish between the intra and extracellular sodium, but can only record the overall TSC, sensitive to both altered volume fractions or altered intra and extra cellular concentrations. Moreover, the spatial resolution of sodium imaging at 3T is poor (with a nominal resolution of 4×4×4mm^3^), which implies that our measurements are currently affected by partial volume. In particular, head motion could affect voxels adjacent to CSF, and be responsible for increase or decrease ΔTSC in such areas. However, acquisition of a sufficient number of (control - rest) images should provide statistical power to account for any motion not sufficiently corrected by registration. Furthermore, the functional paradigm here is non-standard and would benefit from faster image acquisition methods to increase the temporal resolution of the experiment. Nevertheless, the results presented here are all statistically significant and FWE corrected (p<0.001, 20 voxels).

Ultimately, one could speculate on the tissue composition of an imaging voxel further and try to assess the potential contribution to ∆TSC coming from many compartments, defining e.g. fractions of the axonal volume, soma, myelin, extra cellular matrix (including astrocytes), VRSV (or CSF) space and CBV. A comprehensive model should consider exchange of 23Na at synapsis and in other cells such as glial cells, be adapted for different brain regions and different functional tasks (Alahmadi et al., 2017).

This is therefore a proposal for a novel framework, essential for advancing our understanding of the human brain function, where knowledge must bridge gaps between cellular and large-scale systems (D’Angelo and Wheeler-Kingshott, 2017). Dedicated major efforts should be employed to speed up acquisition and at the same time improve spatial resolution of sodium imaging. Potentially, this emerging and exciting field of research could greatly benefit from higher field strength systems (e.g. 7T) (Ranjeva et al., 2018; Riemer et al., 2015). In order to disentangle the sources of the quantitative ∆TSC from fNaI, it would be important to design multi-modal studies that assess a number of variables, such as CBV, CBF, oxygen consumption rates and metabolism for a better estimate of brain energy dynamics (Germuska et al., 2018). Models of fNaI changes could be validated using a range of hemodynamic and physiological signal recordings, including e.g. magnetoencephalography (MEG), near infrared spectroscopy (NIRS) and positron emission tomography (PET), (Shibasaki, 2008).

In conclusion, sodium changes during activity are sufficiently large to be detected and quantified using fNaI *in vivo*. Preliminary quantitative data show encouraging results in terms of coherence of the ∆TSC values between cerebellum and M1 (0.54 vs 0.46 mmol/l). Interpretations of the reduced value of ΔTSC in the CC (0.27 mmol/l) and of the negative ΔTSC in the ipsilateral M1 (−11 mmol/l) (Hamzei et al., 2002) must be cautious and deferred to future studies. Improvements in data acquisition and computational modelling of neurovascular coupling in relation to 23Na flux during action potential generation and maintenance could open a new way forward to assess neuronal activation in humans *in vivo* non-invasively.

## Nomenclature

BOLD: blood oxygenation level dependent
fMRI: functional Magnetic Resonance Imaging
fNaI: functional sodium imaging
MRI: magnetic resonance imaging
TSC: total sodium concentration
ATP: adenosine triphosphate
23Na: sodium ions
GM: gray matter
WM: white matter
M1: primary motor cortex

## Conflict of Interest

The authors declare that the research was conducted in the absence of any commercial or financial relationships that could be construed as a potential conflict of interest.

## Author Contributions

CGWK developed the idea of fNaI; FR contributed to the implementation and data acquisition and analysis; FP contributed to discussion on feasibility; AR helped with data acquisition; GC contributed to image analysis; XG participated to the development of sodium imaging and useful discussion; FP contributed to data reconstruction and analysis; BS contributed to image acquisition and development; EDA supported the idea development and contributed with physiological interpretation.

## Funding and acknowledgements

The NMR unit where this work was performed is supported by grants from the UK Multiple Sclerosis Society and is supported by the UCL/UCLH NIHR (National Institute for Health Research) BRC (Biomedical Research Centre). CGWK also receives funding from the Horizon2020 EU programme (H2020-EU.3.1 (634541)), ISRT, Wings for Life, CHNF. GC has an ECTRIMS fellowship. FPr was funded for the duration of this study by the Medical Research Council (MRC). FPr has a non-Clinical Postdoctoral Guarantors of Brain fellowship. FPa is supported by the Italian Ministry of Health (NET2013-02355313). EDA receives grants from the Human Brain Project, Centro Fermi and the Italian Ministry of Health.

